# Stage-specific ISG expression reveals functional convergence of type I and II IFNs in SIV infection

**DOI:** 10.1101/192021

**Authors:** Nadia Echebli, Nicolas Tchitchek, Stéphanie Dupuy, Timothée Bruel, Caroline Peireira Bittencourt Passaes, Nathalie Bosquet, Roger Le Grand, Christine Bourgeois, Benoit Favier, Rémi Cheynier, Olivier Lambotte, Bruno Vaslin

**Affiliations:** CEA, Université Paris Sud, INSERM U1184, Immunology of Viral Infections and Autoimmune Diseases (IMVA), IDMIT Department / IBFJ, 92265 Fontenay-aux-Roses, France.; Cytokines and Viral Infections, Immunology Infection and Inflammation Department, Institut Cochin, INSERM U1016, Paris, France; CNRS, UMR8104, 75014 Paris, France; Université Paris Descartes, Paris, France.; APHP, Service de Médecine Interne–Immunologie Clinique, Hôpitaux Universitaires Paris Sud, 94270 Le Kremlin-Bicêtre, France.

**Keywords:** SIV, HIV, Type I and type II Interferons (IFNs), Interferon Induced Genes (ISGs), Pathogenesis

## Abstract

Interferons play a major role in controlling viral infections including HIV/SIV infections. Persistent up-regulation of interferon-stimulated-genes (ISGs) is associated with chronic immune activation and progression in SIV/HIV infections, but the respective contribution of different IFNs is unclear. We analyzed the expression of annotated IFN-induced genes in SIV-infected macaques to decrypt the respective roles of type-I (α,β) and type-II (γ) IFNs. Both IFN types were induced in lymph nodes during early stage of primary infection. Induction of type-II IFN persisted during the chronic phase, in contrast to undetectable induction of type-I IFN. Interferome-based analysis of ISGs revealed that at both acute and chronic infection phases most differentially expressed ISGs were inducible by both type-I and type-II IFNs and displayed the highest increases, indicating strong convergence and synergy between type-I and type-II IFNs. The analysis of functional signatures of ISG expression revealed temporal changes in IFN expression patterns identifying phase-specific ISGs. These results suggest that IFN-γ strongly contribute to shape ISG upregulation in addition to type-I IFN and may contribute to progression.

## Introduction

Interferons (IFNs) are among the earliest signaling proteins released by the immune system in response to viral infections (Pestka, 2007; Pestka et al., 2004; Schneider et al., 2014). IFN signaling through IFN receptors ultimately results in transcriptional activation of genes called interferon-stimulated genes (ISGs) (Decker et al., 2002; Der et al., 1998; Rehermann, 2013; Sarasin-Filipowicz et al., 2008), which contribute to the induction of a broad antiviral state against a wide range of pathogens in host cells, limiting viral replication and viral spread (Hubbard et al., 2012; Isaacs and Lindenmann, 1957; Samuel, 2001). ISGs restrict viral infection by blocking key steps of viral replication, inducing the death of infected cells (Chawla-Sarkar et al., 2003; de Veer et al., 2001; Kane et al., 2016; Schneider et al., 2014). They include important modulators of both innate and adaptive immunity (Chang and Altfeld, 2010; Decker et al., 2002; Gonzalez-Navajas et al., 2012; Heim and Thimme, 2014). ISGs act directly on immune cells, favor their recruitment to inflamed tissues, and contribute to inflammation (Gonzalez-Navajas et al., 2012; Rauch et al., 2013; Schroder et al., 2004).

Type I IFN (IFN-α, -β, -ω and type II IFN (IFN-γ) play important roles in HIV/SIV pathogenesis. Type I IFN genes are in the first line of defense against viral infections, including HIV/SIV (Li et al., 2009; Paludan, 2016; Sandler et al., 2014; Schoggins and Rice, 2011). Plasmacytoid dendritic cells (pDC) sense RNA viruses, including HIV (Beignon et al., 2005; Lepelley et al., 2011), and are major contributors of IFN-α to combat acute SIV infections (Bruel et al., 2014; Harris et al., 2010; O’Brien et al., 2013).

IFN-γ, the unique type II IFN (Pestka et al., 2004), is mostly produced by activated NK and T cells, participates in the control of acute infections caused by a variety of viruses, limits pathologies associated with viral persistence, and contributes to adaptive immunity and immune regulation (Pestka et al., 2004; Schroder et al., 2004). The function of IFN-γ in HIV/SIV infections is complex and not completely understood, but chronic activation of HIV-specific T cells results in abundant secretion of this polypeptide in HIV/SIV infections.

Persistent induction of ISG expression in chronic HIV/SIV infections is predictive of progression to pathogenesis (Bosinger et al., 2009; Jacquelin et al., 2009; Lederer et al., 2009), despite their large array of antiviral activities, and IFNs are suspected to drive pathogenesis *in vivo*. Chronic induction of ISG expression is often attributed to type I IFNs, but they are often below the threshold of detection in the asymptomatic chronic phase of infection (Abel et al., 2002; Bosinger et al., 2009; Jacquelin et al., 2009; Malleret et al., 2008). The respective contribution of type I IFNs and IFN-*γ* in the expression of ISGs during chronic infection is still unclear. Although responses to type I IFNs at the early phases of HIV/SIV infection may limit viral replication (Neil and Bieniasz, 2009; Sandler et al., 2014; Stacey et al., 2009), IFN-associated viral signatures and resistance to type I IFNs have been reported (Fenton-May et al., 2013). Most recent studies suggest that type I IFNs may be friends early in infection but foes during the chronic phase (Doyle et al., 2015; Sandler et al., 2014; Zhen et al., 2017). IFN-α expression in acute SIV infection boosts thymic export, but long-term cytokine production leads to premature thymic involution (Dutrieux et al. AIDS, 2014). In addition, acute IFN-α production during primary infection correlates with the activity of indoleamine 2-3-dioxygenase, an immunosuppressive enzyme (Malleret et al., 2008). IFN-γ is also associated with HIV pathogenesis (Roff et al., 2014). Expression levels of IFN-γ-Induced Protein IP10 (CXCL10) is predictive of progression in HIV/SIV infections (Liovat et al., 2012; Ploquin et al., 2016; Roberts et al., 2010).

The investigation of differences in canonical signaling pathways may offer an informative approach to explore specific transcriptomic signatures of type I IFNs and IFN-*γ*. Type I IFNs bind to the heterodimeric transmembrane receptor IFNAR (IFNAR1 and IFNAR2) (Pestka et al., 2004). In the canonical signaling pathway of type I IFNs, IFNAR engagement activates JAK1 and TYK2 receptor-associated protein tyrosine kinases, leading to phosphorylation of the latent cytoplasmic signal transducers and activators of transcription (STAT)1 and STAT2, which dimerize and translocate to the nucleus. They assemble with IFN-regulatory factor 9 (IRF9) to form the trimolecular complex called IFN-stimulated gene factor 3 (ISGF3) (Blaszczyk et al., 2015; Fink and Grandvaux, 2013), which binds to consensus DNA sequences known as IFN-stimulated response elements (ISRE) in gene promoters, activating the transcription of ISGs. IFN-γ signals through the IFN-γ receptor (IFNGR), a heterotetramer of IFN-γ–R1 and IFN-γ-R2 subunits (Bach et al., 1997; Wang et al., 2017). Binding triggers the activation of receptor-associated JAK-1 and JAK-2, and subsequent tyrosine phosphorylation of the cytoplasmic tail of the IFN-γ–R1 subunits. STAT-1 is recruited to the phosphorylated IFN-γR1, becomes phosphorylated, forms homodimers which translocate to the nucleus (Majoros et al., 2017), and binds to gamma-activated sequence elements (GAS) within the promoters of more than 250 IFN-γ-responsive genes (Lackmann et al., 1998).

In this study, we aimed to decipher the respective contributions of type I and type II IFNs to shed light on their respective influence in HIV/SIV infection and pathogenesis. We analyzed IFN and ISG transcriptomic profiles and their associated functional signatures in the cynomolgus macaque (*macaca fascicularis*) pathogenic model of SIV infection, taking advantage of longitudinal access to blood and lymphoid tissues. We elucidated the respective contribution of type I and type II IFNs in determining ISG expression levels, patterns, and associated functions in both acute and chronic SIV infection, by classifying the ISGs into either type I or type II, based on information obtained from the Interferome database (Samarajiwa et al., 2009) or by using an innovative approach to classify ISGs based on the presence of ISRE or GAS conserved consensus sequences in their respective promoters.

## Results

### Viral replication and disease outcome

We investigated the transcriptomic profiles of type I and II IFNs during acute and chronic SIV infection by infecting six cynomolgus macaques with SIVmac251 and longitudinal sampling post infection (p.i.). Plasma viremia peaked on day 10 p.i. (**Figure 1A**) at a mean of 4.4x10^7^ vRNA copies/mL (4.2x10^7^ to 9x10^7^). Five macaques displayed persistently high plasma viral loads during the chronic phase and a decline in the number of CD4 T cells (**Figure 1B**). Animal #30742 had plasma viral loads below 400 copies/mL during the chronic phase and no significant decline in the number of CD4 T cells, thus behaving as a controller. Viral replication remained persistently high during the chronic phase in 5/6 animals in both tissues analyzed (RM biopsies – RB and PLNs) (**Figures 1C and 1D**). Viral RNA loads in tissues were not significantly different at D9 and M3 in these macaques, confirming persistently high SIV replication in tissues during the chronic phase. The vRNA load decreased substantially in animal #30742 tissues by M3, even becoming undetectable in the gut, confirming its controller status.

**Figure 1:**
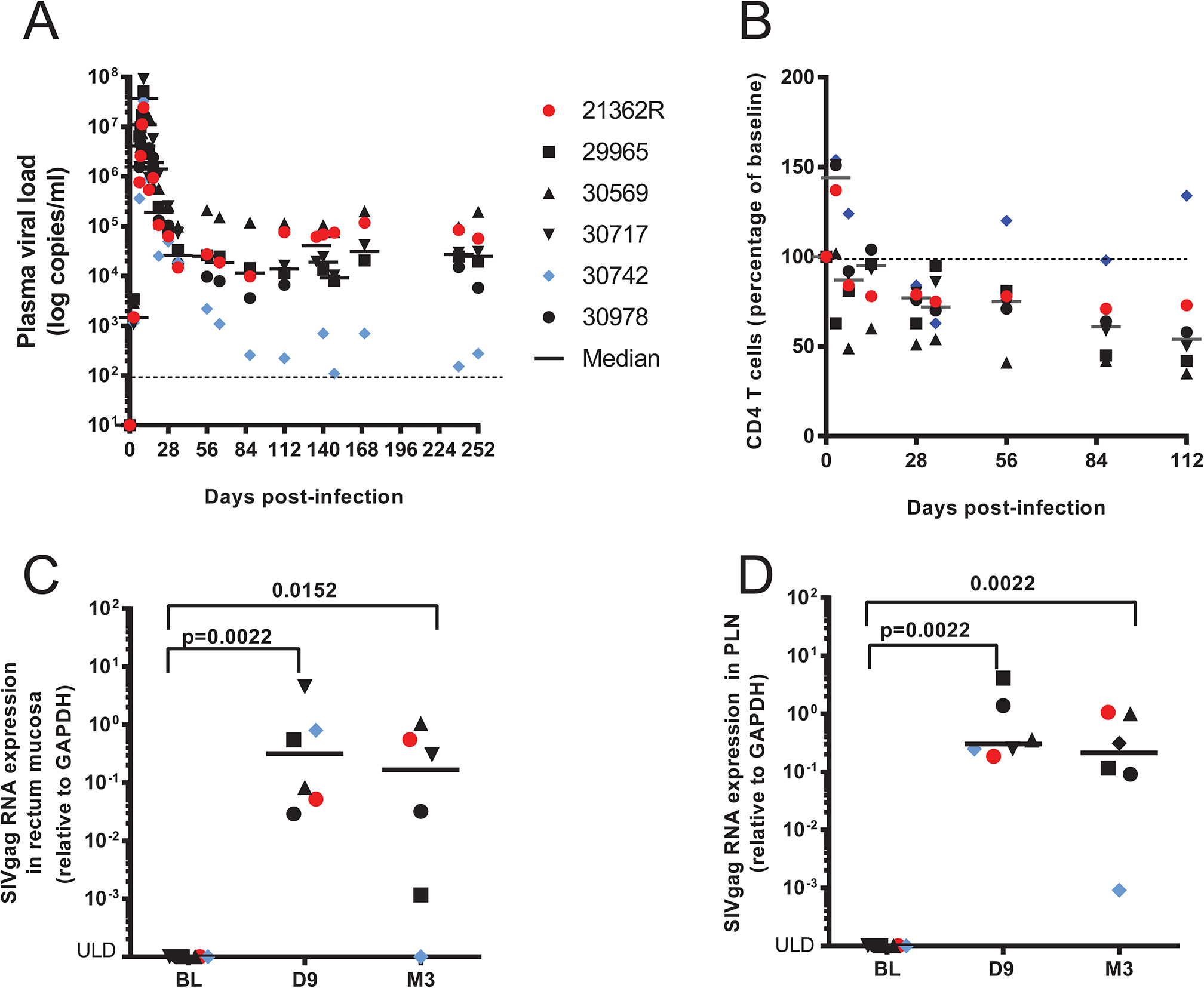
Infection of six cynomolgus macaques with SIVmac251 and establishment of chronic infection. (**A**) vRNA load in plasma. (**B**) Evolution of central memory CD4 T cell numbers in the blood over time (percentage of baseline). (**C**) vRNA in RM. (**D**) vRNA in PLNs.

### Transcriptomic profiling of PBMC, PLN, and RB samples

Before analyzing differentially expressed interferon-induced genes by annotating them using two approaches as described in **Expanded View Figure 1**, we first examined global gene expression profiles in PBMC, PLN, and RB samples longitudinally collected from these animals, before infection and at D9 and M3 p.i., to characterize the molecular signature of SIV infection. We identified genes with significant differential expression from baseline.

Of 43,604 probes, 1,638 genes were upregulated by more than two-fold in at least one condition. Remarkably, we identified many more differentially expressed genes in PLNs and PBMCs than in RBs (**Figure 2A**). Multidimensional scaling revealed strong segregation of the biological conditions (time points and tissues), reflecting large biological differences, as well as the good quality of the gene signatures (**Figure 2B**). We observed the highest segregation between time points for PLNs. More genes were differentially upregulated in PLNs than in RBs or PBMCs, and more genes were differentially expressed at D9 than at M3 in all tissues (**Figure 2C**). Functional enrichment analysis revealed that upregulated genes were mainly associated with functions related to infection, immunity, inflammation, canonical pathways related to interferon signaling, and communication between innate and adaptive immune cells at both phases of infection (**Expanded View Figure 2 and Expanded View Table 1**).

**Figure 2:**
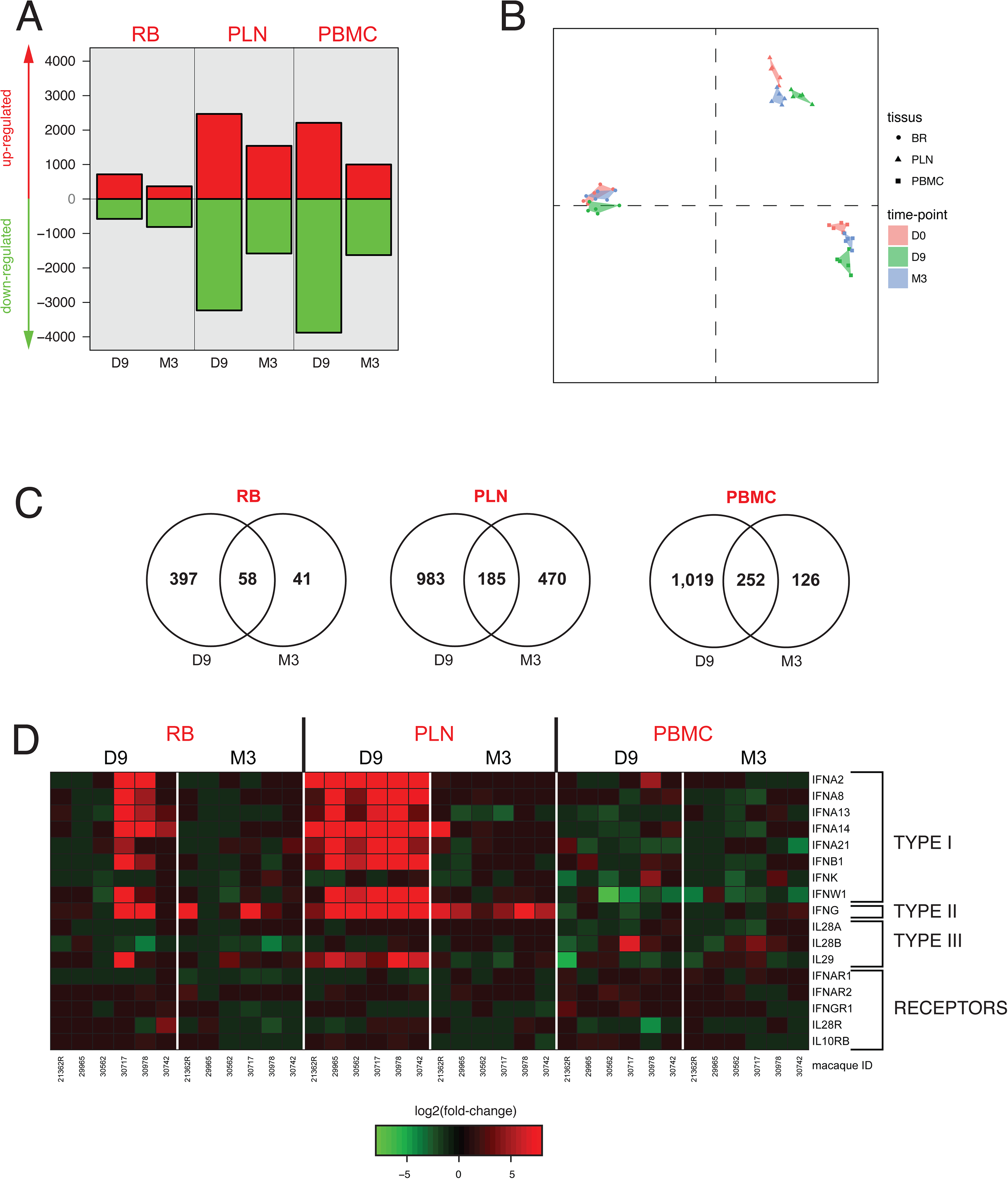
Transcriptome profiling reveals inter-tissue differences and the evolution of differential gene expression post-infection. Two critical time points, day 9 (D9) and month 3 (M3) p.i., were compared to baseline (before viral infection) (**A**) Number of genes with significant differential expression relative to baseline in RBs, PLNs, and PBMCs. (**B**) Multidimensional scaling (MDS) representation showing the segregation of biological samples based on the altered expression of the 1,638 genes with FC > 2. (**C**) Venn diagram showing the distribution of differentially expressed genes over time in RBs, PLNs, and PBMCs. (**D**) Evolution of RNA expression of interferons and interferon receptors in RBs, PLNs, and PBMCs at D9 and M3 p.i. Heatmap representing the FC from baseline for the six SIVmac251 infected macaques.

We first analyzed the expression profile of IFNs and their related receptors, as our aim was to elucidate the respective contribution of type I IFNs and IFN-γ in ISG induction during acute and chronic SIV infection and their contribution to HIV/SIV pathogenesis. There was higher expression of both type I IFN genes (*IFN-α*, *IFN-β* and *IFN-ω*) and IFN-γ during primary than chronic infection. The expression of type I IFN genes was mostly similar in PLNs, whereas it was very heterogeneous in the RBs among the macaques (differential expression in only two of six macaques at D9). No change in type I IFN gene expression was observed in PBMC (**Figure 2D**). *IFN-γ* displayed a similar expression pattern as type I IFN genes during the acute phase, although at lower levels, with similar inter-individual disparity in the RM, and no increase in PBMCs. In contrast to type I IFN genes, high *IFN-γ* expression was remarkably persistent in the PLNs during the chronic phase of infection, likely reflecting ongoing persistent chronic T-cell activation reported in this model (Bruel et al., 2014).

We confirmed the transient increase of *IFN-α* and *IFN-β* mRNA expression during the acute phase of infection by RT-qPCR (**Figure 3A and 3B**). The large inter-individual difference in type I IFN gene expression levels in RBs was also confirmed (**Figure 3C**). We also confirmed the increased expression of *IFN-γ* at both D9 and M3 by RT-qPCR (**Figure 3D**). Microarrays also showed increased ISG expression (**Expanded View Figure 3**) at both time points, confirmed in PLNs by RT-qPCR for *MX1* (**Figure 3E**) and *IRF9* (**Figure 3F**). These data also suggested more consistent ISG expression in PLNs than in RM, likely resulting from the disparity of IFN gene expression in this tissue, and, to a lesser extent, than in PBMCs, in which the expression of IFN genes was not significant at any phase of infection. Acute expression of both type I IFNs and IFN-γ during the primary infection phase was also confirmed at the protein level by measuring the antiviral function of type I IFNs (**Figure 3G**) and IFN-γ concentrations in plasma (**Figure 3H**). During the acute phase, IFN-γ expression, which is inducible by type I IFNs, strongly correlated with that of type I IFNs in the plasma (Pearson R= 0.99, p-value < 0.0001).

**Figure 3:**
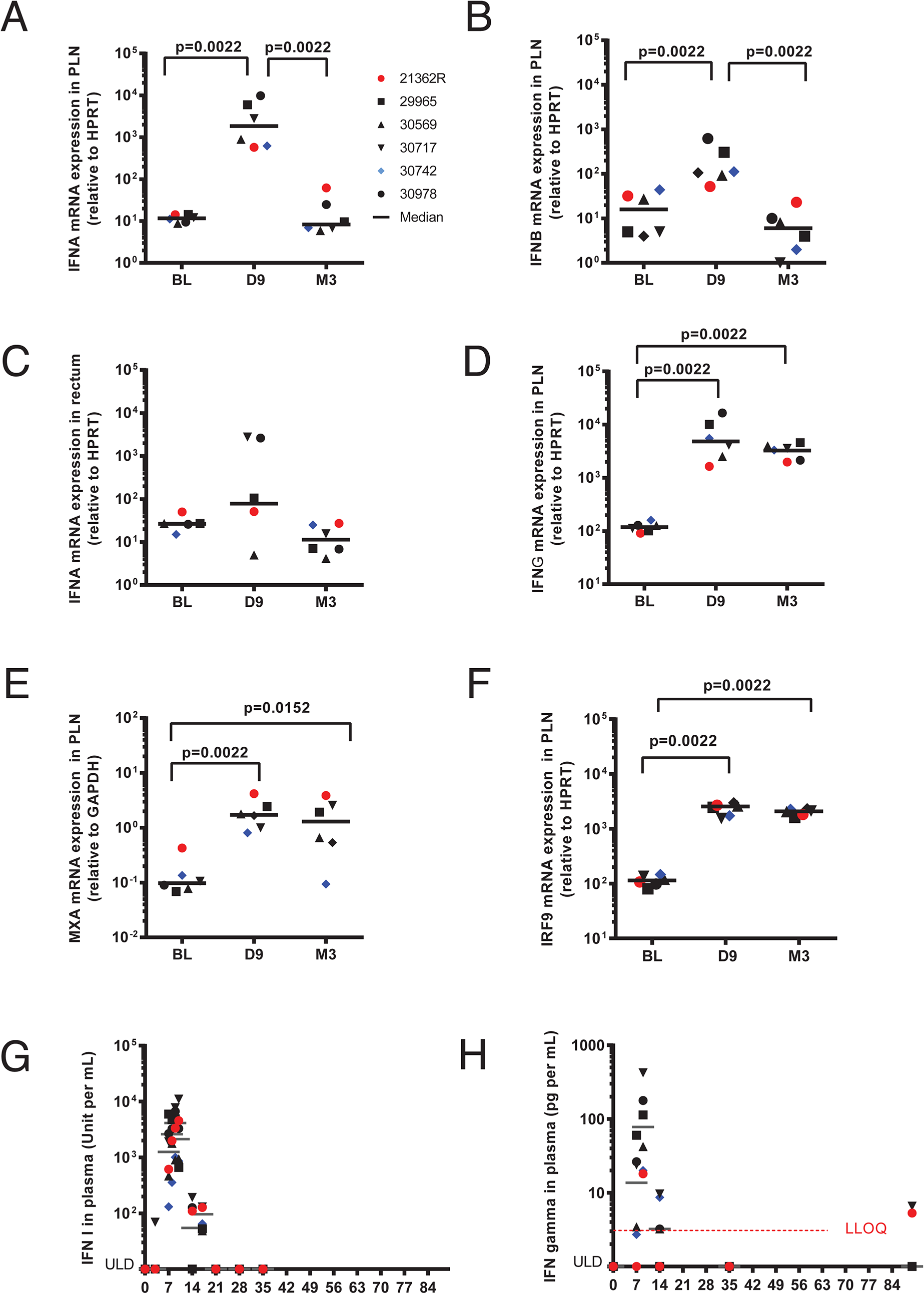
Confirmation of differential expression of IFNs in tissues by RT-qPCR and at the protein level in plasma. (**A**) IFN-α mRNA expression in PLNs by RT-qPCR. (**B**) IFN-β mRNA expression in PLNs by RTqPCR. (**C**) IFN-α mRNA expression in RM by RT-qPCR.(**D**) IFN-γ mRNA expression in PLNs by RT-qPCR. (**E**) MXA mRNA expression in PLNs by RT-qPCR. (**F**) IRF9 mRNA expression in PLNs by RT-qPCR. (**G**) Type-I IFN antiviral activity measured in plasma (MDBK/VSV biological assay) and (**H**) plasma IFN-γ concentration (measured by Luminex).

Our data indicate that PLNs are a major source of IFN production and consequent ISG expression in SIV-infected cynomolgus macaques, in accordance with data reported for SIVmac251-infected rhesus macaques (Abel et al., 2004; Abel et al., 2005; Harris et al., 2010). Persistent IFN-γ expression contrasted with blunted type I IFN gene expression in PLNs during the chronic phase.

### SIVmac251 infection induces both type I and Type II ISGs during both phases of infection in PLNs

We focused our ISG analysis on PLNs, because we previously observed that IFN-α production is followed by high ISG expression in this tissue, and because IFNs were mainly produced in PLNs in the present study. In addition, we previously showed early trafficking of pDC and their contribution to massive IFNα production in PLNs during acute infection (Bruel et al., 2014; Malleret et al., 2008). We excluded the controller macaque #30742 from the data set for ISG analyses to avoid a confounding factor by including an animal with a different disease outcome.

We identified 278 ISGs among 1,638 up-regulated genes in PLNs. A total of 169 ISGs were specifically induced during the acute phase, but not the chronic phase (D9-ISGs), and 58 were specifically induced during the chronic phase, but not the acute phase (M3-ISGs), whereas 51 were induced at both time points (D9&M3-ISGs) (**Figure 4A**). The transition from the acute to chronic phase of infection was associated with a large decrease in the expression of ISGs of the D9&M3-ISG group (p < 0.0001, Wilcoxon signed rank test) (**Figure 4B, left panel**). These genes included apoptotic, activation marker, chemokine/chemokine receptor, interferon regulation, and antiviral genes (**Figure 4B, right panel**). Nevertheless, the change in expression of D9-ISGs was not significantly different from that of M3-ISGs (data not shown). The main biological functions associated with D9-ISGs and M3-ISGs and their FC are listed in **Expanded View Table 2**.

**Figure 4:**
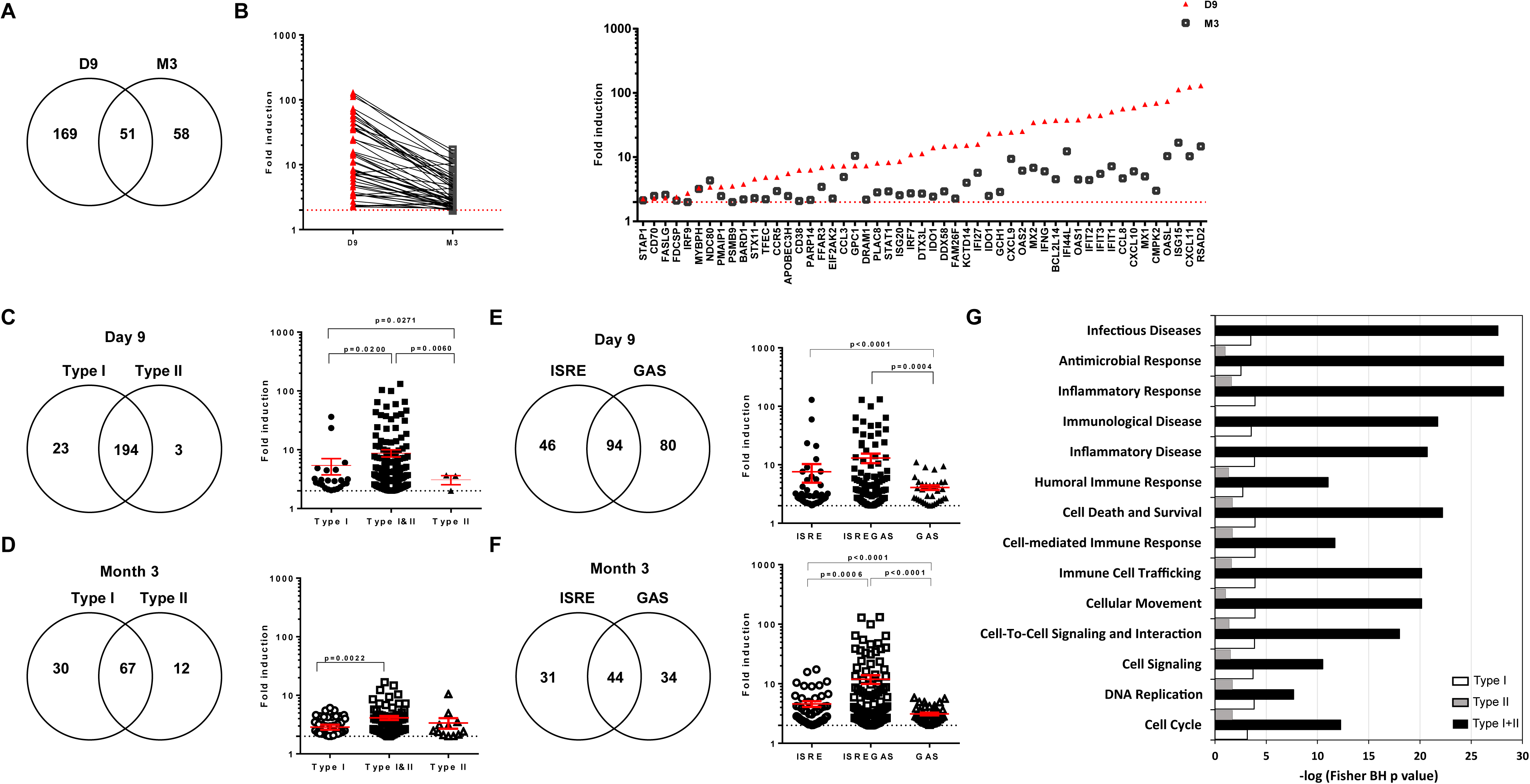
SIV infection induces ISG expression during both phases of infection in PLNs. (**A**) Venn diagram showing the number of ISGs expressed at D9, M3, or both time points. (**B**) Comparison of mean ISG expression FC at D9 and M3 p.i. for the 51 ISGs expressed at both time points. (**C**) Venn diagram of ISGs expressed at D9 p.i., splitting them into those only inducible by type I IFN (Type I), only inducible by type II IFN (Type II), and those inducible by both IFN types (left) and comparison of their mean induction levels (right). (**D**) Venn diagram of ISGs expressed at M3 p.i., splitting them into those only inducible by type I IFN, those only inducible by type II IFN, and those inducible by both IFN types (left) and comparison of their induction levels (right). (**E**) Venn diagram of ISGs expressed at D9 p.i., splitting them into those displaying at least one ISRE, one GAS, or both sequence motifs in their promoters (left) and comparison of their induction levels (right). (**F**) Venn diagram of ISGs expressed at M3 p.i., splitting them into those displaying at least one ISRE, one GAS, or both in their promoters (left) and comparison of their induction levels (right). (**G**) Functional enrichment of type I-, type II-, and type I&II ISG subsets. The p value was calculated by the Fisher test with the Benjamini-Hochberg correction for multiple comparisons. The -log(p-value) is given for the most significant processes. In C, D, E, and F, the p value is given for Welch’s unequal variances *t*-test, considering P < 0.05 to be significant.

We annotated the ISGs using the Interferome database (Samarajiwa et al., 2009) to understand the contribution of type I IFNs, type II IFN, or both to the ISG expression pattern and identified type-I ISGs (inducible by type I IFNs but not type II IFN), type II ISGs (inducible by type II IFN but not type I IFNs), and TypeI&II ISGs (inducible by both IFN species). We also used a novel approach to restrict our analysis to genes directly targeted by interferon signaling and to exclude any genes described in the Interferome database that may be induced by secondary mechanisms. It consisted of annotating the ISGs based on the presence of ISRE and/or GAS consensus sequences in the promoter regions of all ISGs upstream of the 5’UTR, as these sequences are targets of ISGF3 (STAT1/STAT2/IRF9) and STAT1/STAT1 homodimer complexes in the canonical signaling pathways of type I IFNs and type II IFN, respectively.

At D9 p.i. (**Figure 4C, left panel**), 23 genes belonged to the type I ISG group, whereas three belonged to the type II ISG group. These results suggest that type I IFNs and type II IFN induce the expression of distinct and specific sets of ISGs during the acute phase, but most ISGs upregulated at D9 were inducible by both type I IFNs and type II IFN (194 genes, 88.2%), showing strong convergence of type I IFNs and type II IFN in the induction of ISG expression. Type I-ISGs showed a higher FC in expression than type II-ISGs at D9 (**Figure 4C, right panel**) and type I&II-ISGs displayed a significantly higher FC in expression than either type I- or type II-ISGs, suggesting strong synergy between type I IFNs and IFN-γ during the acute phase (p = 0.02 and p = 0.006, respectively).

At M3, 30 genes belonged to the type I-ISG group, whereas 12 belonged to the type II-ISG group (**Figure 4D, left panel**), suggesting that type I IFNs and IFN-γ also induce the expression of distinct sets of ISGs during the chronic phase of infection. Most ISGs significantly induced at M3 were inducible by both type I IFNs and type II IFN (67 genes, 61.5%), as during the acute phase. These ISGs displayed a significantly higher induction of expression than type I-ISGs (p = 0.0022) (**Figure 4D, right panel**). These data show that type I IFNs and IFN-γ strongly converge to induce ISG expression during the chronic phase, and suggest that IFN-γ strongly contributes to enlarging the repertoire of ISGs expressed during the chronic phase.

We then used the GAS/ISRE annotation to investigate the respective roles of the targeting of these sequences in the signaling pathways leading to ISG expression. During primary infection (**Figure 4E**), 94 of 220 stimulated ISGs displayed at least one GAS and one ISRE upstream of the 5’UTR (42.7%). Additionally, 80 displayed at least one GAS, but no ISRE, and 46 displayed at least one ISRE, but no GAS. The expression of ISRE&GAS-inducible ISGs was more highly induced than ISGs inducible through GAS alone (p = 0.0004). These data show that GAS and ISRE signaling pathways induce the expression of distinct ISG subsets (ISRE-inducible ISGs or GAS-inducible ISGs), and converge to induce the expression of GAS&ISRE-inducible ISGs to higher levels than that of ISRE-ISG and GAS-ISG subsets.

In the chronic phase (**Figure 4F**), 31 differentially expressed ISGs belonged to the ISRE-ISG group (33.0%), 34 to the GAS-ISG group (36.2%), and 44 to the ISRE&GAS group (40.4%). The ISRE&GAS-ISGs had, on average, a higher change in expression than the ISRE-ISGs (p = 0.0006) or GAS-ISGs (p < 0.0001). These data show that both GAS and ISRE signaling pathways are used to induce distinct ISG subsets during the chronic phase of SIV infection and that the expression of GAS&ISRE-ISGs is induced to higher levels than that of ISRE-ISGs or GAS-ISGs. This suggests convergence of the type I IFNs and type II IFN canonical pathways and potential synergy between type I IFNs and IFN-γ for the induction of ISG expression.

We then analyzed the biological processes or canonical pathways associated with type-I, type-II, and type-I&II differentially expressed ISG subsets, independently of the time of induction (**Figure 4G**). There was marked enrichment for the three subsets in processes associated with immune responses to infectious disease, including immune-cell trafficking, cell signaling, inflammation, and cell death. Differentially expressed ISGs induced by both type I IFNs and type II IFN (type I&II subset) were much more highly enriched in many immunological functions and inflammation than ISGs only induced by type I IFNs or IFN-γ. These data show that type I&II-ISGs are more highly involved in most of these functions than type-I-ISGs or type-II-ISGs.

Overall, our data show strong convergence of type I IFNs and IFN-γ in the induction of ISG expression in both the acute and chronic phases of SIV infection, as the expression of most ISGs is inducible by both type I IFN and IFN-γ through their respective canonical pathways and show the highest induction.

These ISGs have major functions in inflammation and antiviral responses and represent as much as 88.2% of the upregulated ISGs during the acute phase, when both type I IFNs and IFN-γ are strongly upregulated in PLNs, whereas they represent 61.5% of upregulated ISGs in the chronic phase, when IFN-γ is still upregulated at high levels, but type I IFNs are undetectable. Convergence between type I IFN and IFN-γ was therefore more prominent during the acute than chronic phase, but still played an important role in the induction of ISG expression during the chronic phase.

### The functional signature of ISGs changes during infection

We then analyzed the evolution of the functional signatures of ISGs during infection by comparing their degree of association with biological processes at D9 and M3. The complete list of ISGs, including their FC at each time point and the main functions significantly associated with them, is shown in **Expanded View Table 2**. We then compared three ISG subsets based on their kinetics of induction (ISGs induced only at D9, ISGs only induced at M3, and ISGS induced at both D9 and M3) by Ingenuity Pathway Analysis. The three ISG subsets associated with biological functions with varying strength (**Figure 5**). ISGs induced only at D9 and those induced at both time points were most strongly associated with processes related to cell-mediated immune responses, inflammation, cell death, and the innate immune response to infectious diseases, but weakely associated with DNA replication and cell cycle control, whereas those expressed only at M3 were most significantly associated with two specific functions, cell cycle control and DNA replication. ISGs induced at both time points were most significantly associated with the response to infectious diseases, antimicrobial responses, inflammatory responses, cellular immunity, and cell signaling. These data show that the ISG functional signature is dependent on the time of infection, which was confirmed by comparing signatures of all ISGs induced at D9 to those induced at M3 (**Expanded View Figure 4**).

**Figure 5:**
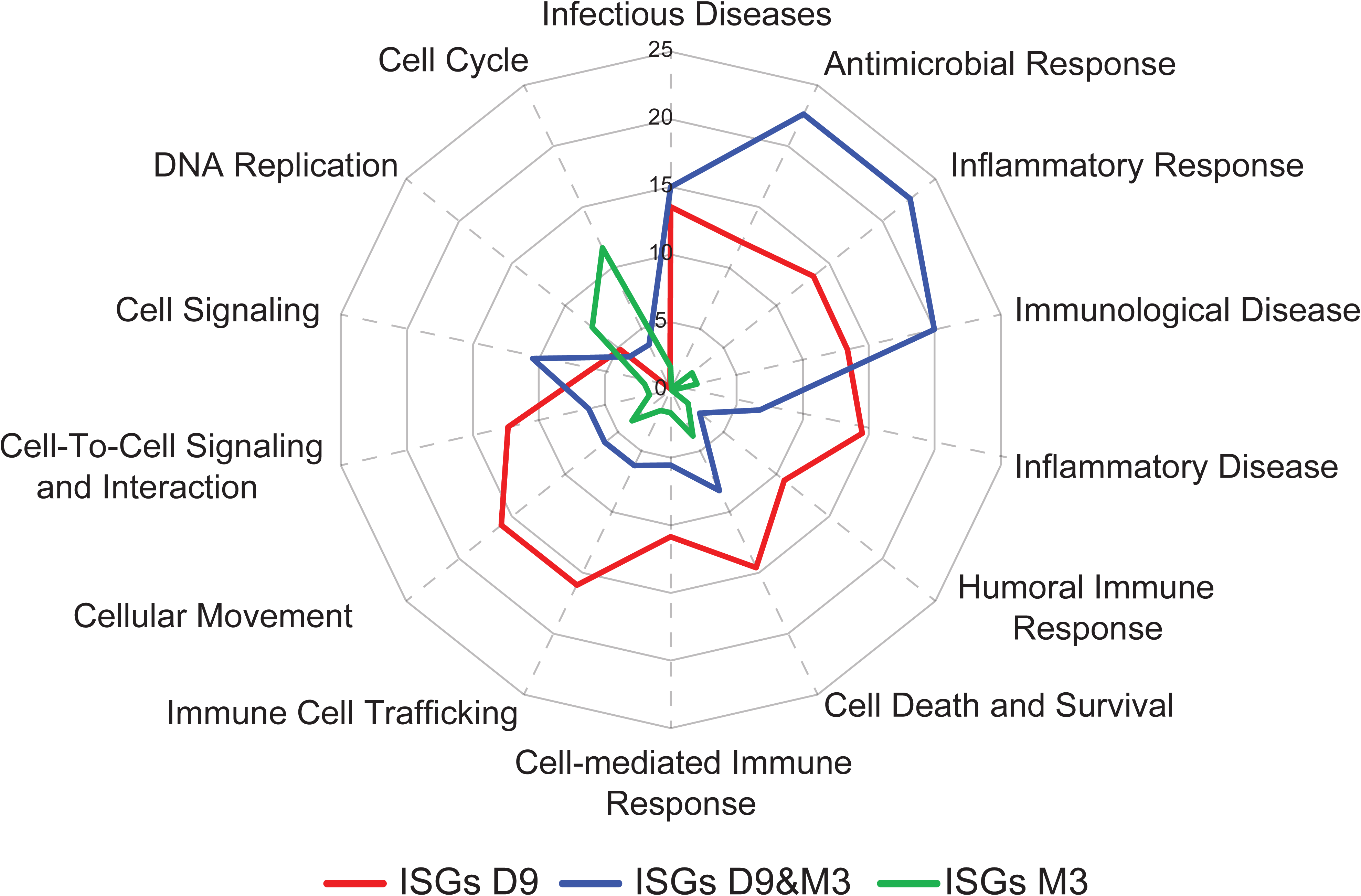
Evolution of the ISG-associated functional signature during infection. Functional enrichment of ISGs differentially expressed at either (**A**) D9 p.i., (**B**) M3 p.i., or (**C**) both time points (D9&M3), was performed using IPA. The right-tailed Fisher Exact Test with the Benjamini-Hochberg correction for multiple tests was used to identify processes showing statistically significant over-representation of focus ISGs. Over-represented functional or pathway processes have more focus genes than expected by chance and the – log(p-values) of the most significant functions are plotted in the radar plot.

We more fully studied the functional imprint of ISGs at the gene level by representing the FC in expression of ISGs that were upregulated and signed the most significant functions in clustered heat maps (**Figure 6)**. They show significant involvement of IFNs in IFN signaling, cell communication (chemokines, cytokines and their receptors), cell activation (both T-cell activation and antigen presenting cells), apoptosis, viral restriction, and cell cycle control, through the induction of ISG expression.

**Figure 6:**
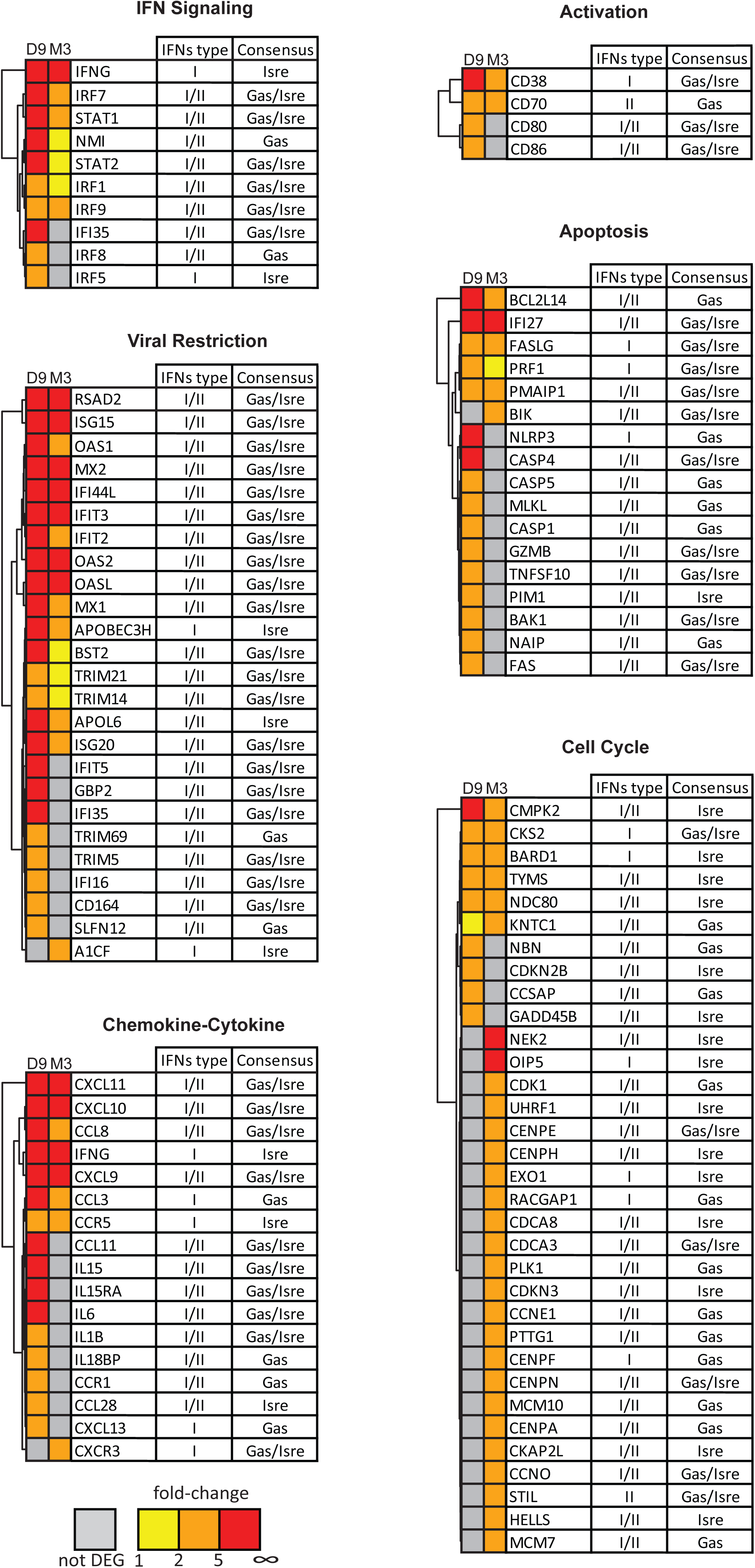
Heat-map signatures of differentially expressed ISGs. The differentially expressed genes of the ISG cluster with the most significant functions are represented as heat-maps. The color intensity shown at each time point is related to the FC in RNA expression: grey for not differentially expressed (not DEG), yellow for FC values between 1 and 2, orange for FC values between 2 and 5, and red for FC values above 5.

IFNs strongly up-regulated the STAT and IRF family transcription factors involved in IFN signaling (**Figure 6-IFN signaling**). Among these, *IFN-γ*, which is an ISG induced by type I IFN, had a mean FC of 36 during the acute phase, which persisted at six-fold in the chronic phase, when induction of type I IFN expression was not detectable. The three components of ISGF3 (STAT1, STAT2, and IRF9) were also strongly induced in the acute phase and persisted during the chronic phase. NMI, which interacts with all STATs, except STAT2, and augments STAT-mediated transcription in response to IFN-γ, was also induced. The persistence of IRF9, which was confirmed by PCR, may enhance the formation and prolonged presence of ISGF3, which would explain the sustained expression of ISRE-ISG, despite low type I IFN expression during the chronic phase. Overall, these IFN signaling genes may contribute to the amplification of IFN production and ISG expression during both the acute and chronic phases.

The most meaningful downstream effect of ISGs induced during both the acute and chronic phase was the antiviral response, including increased expression of well-known antiviral genes: *MX1*, *OAS1*, *OAS2*, *ISG15*, *IFIT* and *TRIM* family genes, and *RSAD2* (tetherin) (**Figure 6-viral restriction)**. Many of these genes are known players in HIV restriction and have antiretroviral activity against HIV/SIV (Abel et al., 2002; Borrow, 2011; Kane et al., 2016; Sandler et al., 2014). *ISG15* and *RSAD2* were the most highly induced (112 and 129-fold in acute infection, respectively). Most of these interferon inducible antiviral genes were strongly induced in the acute phase. Their expression was sustained in the chronic phase, but at a much lower level.

The chemokine/cytokine genes, *CXCL-9*, *-10,* and *-11*, known to be strongly induced by IFN-γ, were induced during both the acute and chronic phases. They likely play an important role in the recruitment of a large variety of immune cells to inflamed lymph nodes at both phases of infection (**Expanded View Figure 4**). The higher expression of *CXCR3*, the common ligand of these CXCL chemokines, during the chronic phase, likely reflects the recruitment of these cells, including activated T cells, and supports the large contribution of IFN-γ in chronic inflammation, as previously suggested in rhesus macaques (Abel et al., 2004). In contrast, the interferon stimulated pro-inflammatory cytokine genes, *IL-6*, *IL-IB*, *IL15*, *IL15-RA,* and anti-inflammatory gene *IL18BP*, were induced only during the acute phase but were not sustained during the chronic phase showing that IFNs may contribute more strongly to inflammation during the acute than chronic phase, while still maintaining cell trafficking in the lymph nodes during the chronic phase through induction by CXCL chemokines. This controlled expression of IFN-induced pro-inflammatory cytokines in the transition from the acute to chronic phase paralleled the reduction of type I IFN expression to undetectable levels and that of type II IFN to chronic levels (lower than acute phase), although we observed persistently high viral replication in PLNs. *CCL3* and *CCR5* expression increased in both phases and may contribute to persistent migration of CCR5**^+^** target cells to the lymph nodes to fuel SIV replication during the chronic phase, sustaining the observed high level of viral replication (**Figure 1D**). Overall, these results show that many ISGs are key players in SIV induced inflammation and cell trafficking in lymph nodes, with notably different patterns during acute and chronic infection, likely due to changes in the ratio of type I IFN to type II IFN expression over time.

Among differentially expressed ISGs, activation markers, both T-cell (*CD38*) and antigen presenting cell markers (*CD70*, *CD80*, *CD86*), were also induced during the acute phase (**Figure 6-activation**), and *CD38* and *CD70* (CD27 ligand) expression persisted during the chronic phase, likely related to the chronic T-cell activation that occurs in this model (Bruel et al., 2014).

Strong IFN production during acute infection was also associated with increased expression of numerous pro-apoptotic ISGs (**Figure 6-apoptosis**). Most were strongly induced during the acute phase, but their expression during the chronic phase was much lower and restricted to a few genes. These data suggest that IFNs strongly contribute to the regulation of apoptosis in PLNs both during both primary infection and the chronic phase.

The heat map for cell cycle-ISGs (**Figure 6-cell cycle**) show that the acute phase of SIV infection, which was associated with high type I IFN expression in PLN, was associated with increased expression of anti-proliferative ISGs, such as *GADD45B*, *CDKN2B*, and *NBN*, which act at different points to block the cell cycle. In contrast, ISGs induced exclusively during the chronic phase were most strongly associated with processes related to the G1 phase, genes involved in DNA replication and control, and G1/S and G2/M phase transitions. This set of ISGs included *CDK1*, a cyclin-dependent kinase, *CCNE1*, *MCM7*, *NEK2*, *CDCA8*, *CENPE*, *CENPH*, and *CENPA*. This expression profile, including expression of these cyclins and CDK genes, was specific to the chronic phase, and thus associated with persistent IFN-γ expression and undetectable type I IFN expression, suggesting a positive role of IFN-γ on the cell cycle.

## Discussion

Chronic expression of ISGs is characteristic of pathogenic SIV/HIV infections (Bosinger et al., 2009; Jacquelin et al., 2009; Lederer et al., 2009), but the respective contribution of IFN species to the induction of ISG expression is unclear.

The aim of the present study was to discriminate the role of type I and type II IFNs in shaping the pattern of ISG expression over time, and to study the dynamics of biological functions downstream of ISG expression during the acute-to-chronic transition during SIV infection to better understand their role in pathogenesis and immune regulation. We performed a comprehensive analysis of differentially expressed ISGs by DNA array in PLNs of SIV infected monkeys. We considered the level of expression (relative to pre-infection levels), the ability to be induced by type I and/or type II IFNs, and the existence of IFN-signaling motifs in the promoter (ISRE and GAS) of each differentially expressed ISG to investigate the signaling pathways involved. In parallel, we monitored IFN expression to correlate ISG and IFN expression patterns.

High expression of both type I and type II IFNs in PLNs, during the acute phase of infection, switched to persistent IFN-γ expression and type I IFN expression below the threshold of detection during the chronic phase. This observation is in accordance with other models of pathogenic SIV infection (Abel et al., 2005; Khatissian et al., 1996). Our results clearly show that *IFN-α*, *IFN-β*, *IFN-ω* and *IFN-γ* were strongly expressed in PLNs, to a lesser extent in RM (with high inter-individual variation), and not in PBMCs, in accordance with the finding of Abel *et al.* (Abel et al., 2002; Abel et al., 2004). These data highlight lymph nodes as a major source of IFNs during primary infection. Our data contrast with previously published microarray studies in which type I IFNs were not detected by DNA arrays in either blood, PBMCs, blood CD4 T cells, or PLN cells in pathogenic models (Bosinger et al., 2009; Jacquelin et al., 2009; Lederer et al., 2009). This may be due to differences in the time of sampling after infection, sensitivity of the microarrays used, or the type of tissue analyzed. Indeed, our choice of sampling PLNs at the peak of type I IFN levels in plasma, based on our knowledge of the model and the early cytokine burst in HIV-infection (Bruel et al., 2014; Stacey et al., 2009), was critical, as type I IFN was only consistently detectable in PLNs during the early phase of primary infection. Use of the Agilent macaque DNA array platform, which offers both a high dynamic range of gene expression measurements and high sensitivity, was also likely critical for measuring the expression of type I IFN genes. Indeed, microarray and RT-qPCR measurements in PLN were positively correlated (p < 0.0001, r = 0.9881 for *IFN-α*; p = 0.0003, r = 0.8639 for IFN-γ).

Changes of IFN expression patterns over time were associated with the evolution of the ISG expression pattern. During the acute phase of infection, we observed 220 differentially expressed ISGs, in accordance with pioneering work in other pathogenic models (Bosinger et al., 2009; Jacquelin et al., 2009; Lederer et al., 2009), which showed strong IFN responses to primary SIV infections. Our work is original because we concentrated our analysis on the origin of ISG expression. We identified three classes of ISGs, type I-ISGs (inducible by type I, but not type II IFNs), type II-ISGs (inducible by type II, but not type I IFNs), and type I&II-ISGs (inducible by both type I and type II IFNs), which were the most abundant. Fewer ISGs were expressed during the chronic phase (109 genes) than in the acute phase (220 genes) of infection, but we observed a similar distribution pattern for these three classes at the latter time point (type I&II-ISGs > type I-ISGs > type II-ISGs). The transition from acute-to-chronic infection was also associated with changes in the functional pattern of differentially expressed ISGs, and a decrease of their expression. Indeed, ISGs induced during the acute phase were either expressed at lower levels during the chronic phase, or not at all. However, there was a specific group of ISGs, which were not differentially expressed during the acute phase, for which the expression increased during the chronic phase. This specific ISG set was mostly composed of genes involved in cell cycle control and mitosis, likely important for continuous cell division in inflamed lymph nodes. These data support the conclusion that both type I IFNs and IFN-γ specifically imprint the ISG expression pattern during both phases of infection (type I-ISGs and type II-ISGs, respectively) and converge and synergize for the induction of a large common set of ISGs (type I&II-ISGs) that exhibit major antiviral immune functions. Our results also suggest that convergence of type I and type II IFNs is important during both phases of pathogenic SIV infection for sustained expression of the large set of type I&II ISGs at higher levels than either type I or type II-ISGs, which also show statistically more restricted expression (both in pattern size and intensity). These data suggest that persistent expression of a large array of ISGs during the chronic phase could be largely due to IFN-γω fueled by chronic T-cell activation (Cha et al., 2014; Rehermann, 2013; Zuniga et al., 2015), and NK cells (Jacquelin et al., 2014), the main producers of IFN-γ. Indeed, the expression of several well-known IFNγ-inducible ISGs, such as Mig (*CXCL9*) and IP-10 (*CXCL10*) were sustained during the chronic phase in PLNs. The parallel increase in the expression of their receptor, CXCR3, is consistent with a previous report on rhesus macaques, suggesting that IFN-γ induced Mig/*CXCL9* and IP-10/*CXCL10* expression results in the recruitment of CXCR3^+^-activated T cells, promoting SIV replication and persistent inflammation (Abel et al., 2004).

The finding that 13 ISGs, differentially expressed during the chronic phase, harbor an ISRE but no GAS in their promoters (**Expanded View Figure 5**), suggests possible induction by low type I IFN production through the ISGF3/ISRE canonical pathway, although type I IFN was not detectable during the chronic phase. Alternatively, these genes could also be induced by other transcriptional factors (Ivashkiv and Donlin, 2014; Schoggins and Rice, 2011), independently of IFN, or by non-canonical type II IFN signaling. Indeed, several studies demonstrated ISGF3 complex activation following IFN-γ treatment in murine cells (Morrow et al., 2011), an alternative type II IFN signaling pathway targeting ISREs, suggesting a role for ISGF3 in the convergence and synergy of type I and type II IFN signaling. ISGF3 can induce expression of type II IFN, which is a type I inducible gene, thereby further amplifying the response to type I IFN. We observed increased expression of both STAT1 and IRF9 during the acute phase and persistence of their expression during the chronic phase of infection, albeit at lower levels. Type I IFN expression was not detectable during the chronic phase, in contrast to IFN-γ. Thus, these data suggest that type II IFN may contribute to STAT1 and IRF9 expression and the formation and prolonged persistence of ISGF3 to target ISRE harboring ISGs. Indeed, the number of expressed ISRE harboring ISGs, with no GAS, was still high during the chronic phase, despite the transition from high to undetectable type I IFN expression. Based on these observations, we propose that chronic IFN-γ induction could contribute to induce the expression of ISRE harboring genes through ISGF3 signaling. Indeed, in our study, two genes, which are only type II IFN inducible, fit this paradigm, as they were expressed during the chronic phase, suggesting that there is such an alternative pathway *in vivo*, as reported *in vitro* (Morrow et al., 2011). However, their incidence is probably low, as all other type II IFN-inducible ISGs bear GAS (**Expanded View Figure 5)**.

Our data also suggest that alternative type I IFN signaling takes place *in vivo*. Type I IFN can induce canonical type II IFN-signaling molecules through the induction of active STAT1 homodimers, which can bind to GAS sites to activate the transcription of target genes (Platanias, 2005). In our study, 13 and 11 genes differentially expressed during the acute or chronic phase, respectively, fit this paradigm, as they are only inducible by type I IFNs but harbor only GAS and not ISRE in their promoters (**Expanded View Figure 5**).

Type I and type II IFN signaling pathways crosstalk at other levels and can activate distinct and common ISGs that govern their antiviral effects. Most responses to type I and type II IFNs require the STAT1 transcription factor (Beutler et al., 2005). In our model, differential STAT1 expression at both phases of infection could be a possible point of synergy between type I and type II IFNs. STAT1 expression is inducible by both IFN species and harbors both ISRE and GAS in its promoters, making it a central element in type I and type II IFN crosstalk. Our DNA array study in lymphatic tissue also revealed phase-specific IFN-induced gene expression and highlights the role of type I and type II interferons in the antiviral response to SIV infection and their contribution to numerous biological functions. The interferon response plays an important role in innate immunity against viruses (Wong and Chen, 2016). The strong and early IFN signaling observed here is consistent with reports suggesting that a strong innate immune response in lymph nodes is a mechanism for limiting SIV pathogenesis (Pereira et al., 2008). It was previously shown that IFNs can block SIV replication at early phases of infection *in vitro* (Borrow et al., 2010; Taylor et al., 1998). Although type I and type II IFNs likely inhibit viral replication and activate anti-viral immune responses, these responses are not sufficient to prevent chronic infection. The ineffectiveness of the IFN response in containing viral infection could result from insufficient production of most efficient cytokines (Gibbert et al., 2013) or viral resistance to IFNs during infection. Our current analysis indicates that type I IFN synergizes with type II IFN early to induce anti-viral and restriction factors that may help to limit viral replication *in vivo*. Indeed, blocking type I IFN signaling early in infection was recently shown to worsen SIV infection (Cheng et al., 2017). The early IFN response could participate in the selection of type I IFN-resistant variants, as observed in HIV primary infection (Fenton-May et al., 2013). Lower type I IFN expression during the chronic phase, likely responsible for the observed reduced levels and repertoire of restriction factors in our study, may release viral progeny from IFN pressure. Indeed, viral isolates obtained during the chronic phase of HIV infection are less resistant than viruses isolated during primary infection early after the acute IFN response (Fenton-May et al., 2013).

Several differentially expressed ISGs encode cytokines, chemokines, and their receptors, involved in inflammation and cell recruitment to inflamed lymph nodes. Primary infection was characterized by a burst of expression of IFNs and IFN-induced pro-inflammatory cytokines (IL-1b, IL-6, IL-15 and IFN-γ), chemokines (CXCL9,10,11, CCL3, 8, 11, 28), and IFN-induced chemokine receptors (CCR1, CCR5, CXCR3), whereas this pattern was reduced to IFNγ in the chronic phase, with undetectable type I IFN and no other IFN-induced pro-inflammatory cytokines, despite persistent viral replication in the PLNs. Although most of these genes are also inducible by other factors (Beq et al., 2009; Ponte et al., 2017), most are inducible by both type I IFNs and type II IFN. This suggests that high type I IFN and type II IFN expression during primary infection synergize to take part in inflammation in lymph nodes, creating a favorable milieu for immune cell recruitment. During the chronic phase, in a much lower pro-inflammatory context, including undetectable type I IFN, IFN-γ likely remains one of the few major actors of inflammation and cell recruitment in lymph nodes. Indeed, persistent high IFNγ expression was associated with persistent expression of IFNγ inducible CXCR3 ligands, including CXCL9,10,11, which may contribute to persistent recruitment of immune cells, such as pDC, which persistently accumulate in lymph nodes during chronic SIV infection in our model (Bruel et al., 2014; Malleret et al., 2008), express CXCR3 and CCR5 (Penna et al., 2001), and respond to CXCR3 ligands (Krug et al., 2002). Activation of pDC in lymph nodes (Bruel et al., 2014) may also contribute to CCL3 expression, further increasing the attraction of pDCs and other CCR5^+^ cells. Increased expression of CCL3 may be a beneficial response of the host against HIV/SIV infection, due to its ability to block HIV entry into CCR5^+^ T cells (Cocchi et al., 1995), but it can also be deleterious, through the recruitment of new target cells to inflamed lymph nodes. The CCL3 gene is reported to be type I IFN-inducible in the Interferome database. However, it bears a GAS sequence, but no ISRE, in its promoter and could thus be sustained by IFN-γ during the chronic phase or low undetectable type I IFN expression through alternative type I IFN signaling.

The depletion of CD4 T cells is a signature of SIV and HIV pathogenesis that drives disease progression (Deeks, 2011). We observed a significant decline in the number of CD4 T cells below baseline levels. The pathway causing CD4 T-cell death in HIV-infected hosts is poorly understood (Thomas, 2009), but may involve IFN responses. Most (95%) quiescent lymphoid CD4 T cells die by caspase-1-mediated pyroptosis (Eckstein et al., 2001). In contrast to apoptosis, pyroptosis is a highly inflammatory form of programmed cell death. Caspase-1 (Casp1) is the best-described inflammatory caspase. It processes the IL-1β and IL-18 precursors and induces pyroptotic cell death. Our study highlights the significant differential expression of several mediators of pyroptosis, such as IL1-β, Casp1, NLRP3, and IL15 during acute infection and the absence of induction of Casp3 expression. The involvement of IFNs in pyroptosis could therefore contribute to the early depletion of CD4 T cells in lymph nodes and other tissues. These pyroptosis mediators were barely detectable during the chronic phase, when type I IFN was also undetectable, and two of them (Casp1 and NLRP3) are GAS activated genes whereas IL-15 and IL1b are GAS&ISRE ISGs, suggesting IFN-γ may play a more important role than type I IFN. In non-pathogenic SIV infections, Casp3-dependent apoptosis of productively infected cells may be responsible for most cell death rather than that mediated by Casp1, thus avoiding local inflammation (Doitsh et al., 2014).

IFNs are also known for their anti-proliferative effects (Chawla-Sarkar et al., 2003; Zhao et al., 2013). Our data show that during the acute phase of SIV infection, associated with strong expression of both type I and type II IFN in lymph nodes, differentially expressed ISGs involved in the cell cycle included more anti-proliferative genes (CDKN2B, NBN, GAB45B) which act at different phases to block the cell cycle, than during the chronic phase (CDKN3), at a time when type I IFN expression was reduced to undetectable levels and that of type II IFN persisted. Indeed, during the chronic phase, cell cycle-associated ISGs were more strongly associated with functions/mechanisms that support progression through the cell cycle (related to G1 the phase, control of DNA replication, G1/S and G2/M phase transitions). This is in accordance with the reported synergy between IFN-α and IFN-γ to promote anti-proliferative effects (Schiller et al., 1986), and suggests that coordinated expression of IFN-α and IFN-γ (which is inducible by IFN-α) during the peak of IFN-α production during primary infection may transiently delay immune cell expansion. Indeed, lower numbers of proliferative T and B cells were previously found in PLNs at day 8 or 11 than at day15 p.i. in this model (Benlhassan-Chahour et al., 2003), when type I and type II IFN production is already much lower. Our data suggest that when type I IFN falls to undetectable levels, persistent type II IFN production during the chronic phase results in the induction of a specific set of ISGs likely more favorable for the division of immune cells. We previously reported that cycling T cells increased after day 14 p.i. in PLNs and blood in our model (Benlhassan-Chahour et al., 2003), at a time when type I IFN-I had already dropped to low levels (Malleret et al., 2008). This paradigm is also supported by data reported in mice showing that IFN-γ increases the entry of lymph node CD4^+^ T cells into the cell cycle, favoring cell expansion (Reed et al., 2008). Persistent IFN-γ expression could therefore facilitate T-cell cycling through the induction of phase-specific ISGs during the chronic phase, when levels of anti-proliferative ISGs (CDKN2B, NBN, GAB45B), associated with type I IFN expression, have dropped. Overall, our data suggests that the balance between type I and type II IFN levels is important in regulating the cell cycle during the different phases of infection.

In conclusion, this global analysis, focusing on type I and type II ISG expression patterns, revealed that most differentially expressed ISGs are inducible by both type I and type II IFNs. Furthermore, ISG induced by both type I and type II IFNs were the most highly induced, suggesting that both IFN types converge and synergize to promote many biological functions. This phase-specific ISG response likely results from the switch from a type I to type II IFN-dominated ratio and highlights the significance of IFN-γ in shaping ISG response during the chronic phase. Although these genes could also be co-regulated by other transcriptional factors, a potential confounding and limiting factor in our study, the identification of phase-specific IFN-regulated genes advances our knowledge of the complex role of IFN species in HIV/SIV pathogenesis, which may pave the way to improve therapeutics that target specific host or virally-induced IFN pathways.

## Materials and Methods

### Ethics statement

Six adult male cynomolgus macaques (*Macaca fascicularis*) were imported from Mauritius and housed in the facilities of the “Commissariat à l’Energie Atomique et aux Energies Alternatives” (CEA, Fontenay-aux-Roses, France). Non-human primates (NHP, which includes *M. fascicularis*) are used at the CEA in accordance with French national regulations and under the supervision of national veterinary inspectors (CEA Permit Number A 92-032-02). The CEA complies with the Standards for Human Care and Use of Laboratory Animals of the Office for Laboratory Animal Welfare (OLAW, USA) under OLAW Assurance number #A5826-01. All experimental procedures were conducted according to European Directive 2010/63 (recommendation number 9) on the protection of animals used for scientific purposes. The animals were used under the supervision of the veterinarians in charge of the animal facility. This study was accredited under statement number 12–103, by the ethics committee “Comité d’Ethique en Expérimentation Animale du CEA” registered under number 44 by the French Ministry of Research.

### Animal infection and sample collection

Cynomolgus macaques (n = 6) were exposed intravenously to 5,000 AID50 SIVmac251 as previously described (Bruel et al., 2014; Karlsson et al., 2007). Peripheral Blood Mononuclear Cells (PBMCs), Peripheral Lymph Nodes (PLNs), and Rectal Mucosa (RM) were collected longitudinally. cDNA microarray profiling was performed for both acute (day 9 post-infection: D9) and chronic phases (month 3 post-infection: M3) of infection.

### Viral RNA and T-cell quantification

Plasma vRNA was assayed as previously described (Benlhassan-Chahour et al., 2003; Karlsson et al., 2007). Absolute T-cell counts were calculated from lymphocyte counts obtained by automated cell counting (Coulter MDII; Coultronics, Villepinte, France) combined with flow cytometry data as previously described (Bruel et al., 2014).

### IFN-I antiviral activity

Type I IFN activity in plasma was measured using a bioassay that assesses the reduction of the cytopathic effects induced by vesicular stomatitis virus in Madin-Darby bovine kidney cells as previously described (Bruel et al., 2014).

### RNA extraction and qPCR

Total RNA was isolated from tissues using the RNeasy mini kit (Qiagen), according to the manufacturer’s instructions. To avoid genomic DNA contamination, an additional DNAse step was included after the RNAse-free DNAse step (Qiagen), according to the kit recommendations. The quality and quantity of the RNA were determined using a Nanodrop spectrophotometer. The quantitative rev-transcriptase Kit (Qiagen) was used to synthesize cDNA. Quantitative PCR was performed for each sample on a Light-Cycler using SYBERgreen (Roche) and 0.2 µl FastStart Taq (Roche) in a final volume of 25 µl.

Quantitative RT-PCR was used to assay *IFN-γ*, *IRF9*, and *MXA* gene expression using primers described in **Expanded View Table 1**. For *IFN-γ* and *IRF9*, q-PCR was performed using the following program: 5 min at 94°C, 40 times (15s 94°C; 30 s 56°C, and 30s 68°C), and 10min at 68°C. The amplification of *MXA* was performed under the following conditions: 95 °C for 3 min, followed by 50 cycles of (10s 95 °C, 30s 58°C, and 45s 72°C). *HPRT* or *GAPDH* gene expression was used to normalize mRNA levels. Detection of a single product was verified by dissociation curve analysis and relative quantities of mRNA calculated using the method described by (Pfaffl, 2001). Quantitative RT-PCR was used to assay IFN-*α* and IFN-β as described previously (Bruel et al., 2014; Dutrieux et al., 2014).

### Transcriptome profiling

For oligonucleotide array analyses, total RNA was isolated using the RNeasy RNA isolation kit (Qiagen, CA). The quality and quantity of the RNA were determined using a Bioanalyzer 2100 (Agilent technologies). The cRNA was then amplified using the Low Input Quick Amp labeling kit, one color (Agilent technologies). The quantity and quality of cRNA were evaluated by capillary electrophoresis using an Agilent Technologies 2100 Bioanalyzer and NanoDrop ND-1000. Probe labeling and microarray hybridizations were performed as described in the Agilent 60-mer oligo microarray processing protocol (Agilent Technologies, CA). Six hundred ng of each labeled RNA sample was hybridized to Agilent 8X60K rhesus macaque *Macaca Mulatta* custom 8*60K array (AMADID 045743, Agilent technologies). The Agilent One-Color Microarray-Based Gene Expression Analysis Protocol was followed for hybridization and array washing. Arrays were scanned with an Agilent microarray G2505C scanner and image analysis was performed using Agilent Feature Extraction 11.0.1 Software. Raw images were analyzed using Agilent Feature Extraction software (version 9.5.3.1) and the GE1_1100_Jul11 extraction protocol. All arrays were required to pass Agilent QC flags.

### Transcriptomic analysis

Microarray data were processed using R/Bioconductor. Gene expression values were scaled and normalized using the limma package. Differentially expressed genes were identified using two-sample t-tests (p < 0.05) and based on a 2-fold-change (FC) threshold. Functional Enrichment analysis was performed using Ingenuity® Pathway Analysis (IPA, version n° 31813283) and the right tailed-Fisher Exact test. P-values were corrected for multiple comparisons using the Benjamini-Hochberg Multiple Testing Correction. Genes were annotated using DAVID Bioinformatics Resources 6.8 (NIAID/NIH, USA).

Raw and normalized microarray expression data will be publicly available on the ArrayExpress database upon manuscript acceptance (ID: E-MTAB-6068). These data are temporary available at ftp://188.165.13.182/public/.

### ISG annotations

Differentially expressed interferon-induced genes were annotated using two approaches as described in **Expanded View Figure 1**. The first approach was based on the Interferome database (http://www.interferome.org), an open-access database of type I, II, and III Interferon-regulated genes collected from analyzing expression datasets of cells and organisms treated with IFNs. We used this database to identify and classify genes depending on their expression following induction by either type I IFN, type II IFN, or both IFN species. We also used a complementary approach in which the genes were annotated depending on the presence of either GAS, ISRE, or both consensus sequence motifs in gene promoters within the 2,000 bp upstream of the 5’UTR after locating them using Geneious software (http://www.geneious.com/). This analysis aimed to identify ISG sets downstream of the canonical type I IFN signaling pathway (ISRE ISGs), the canonical type II IFN signaling pathway (GAS ISGs), or both (GAS&ISRE ISGs).

## Acknowledgments

The authors thank Hybrigenics-Helixio (former Imaxio) to which the microarrays were sub-contracted for technical advice, quality controls of RNA samples, microarrays, and initial statistical analysis. The authors are particularly thankful to Véronique VIDAL and Valérie CHABAUD from HELIXIO, for their efficiency and technical advices.

## Author contributions

Conceived experiment and supervized the work: B.V.; Performed experiment: N.E., S.D, T.B., C.P.B.P., N.B.; Secured funding: B.V., R.L.G.; Analyzed microarray data N.E., N.T., B.V.; Wrote the manuscript: N.E., N.T., B.V.; Provided expertise, feedback and critical review of the manuscript: R.L.G, R.C., B.F., C.B., O.L., T.B., C.P.B.P.

## Expanded View Figure and Table Legends

**Expanded View Table 1: Functional enrichment analysis of PLNs, RBs, and PBMCs at D9 and M3.** Global functional enrichment analysis was performed on all differentially expressed genes independently of their FC. The highest and lowest –log (p-values) of all significant functions within each category are given for both D9 and M3 p.i. for each tissue, including PLNs (**A**) RBs (**B**), and PBMCs (**C**).

**Expanded View Table 2:** List of ISGs differentially expressed at either D9 **(A)**, M3 **(B)**, or both time points **(C)** with their mean FC and associated functions.

**Expanded View Table 3: primers and probes.** Sequences of the primers and probes used for quantitative RT-PCR analysis. All sequences are shown 5’ to 3’.

---

**EV Figure 1:**
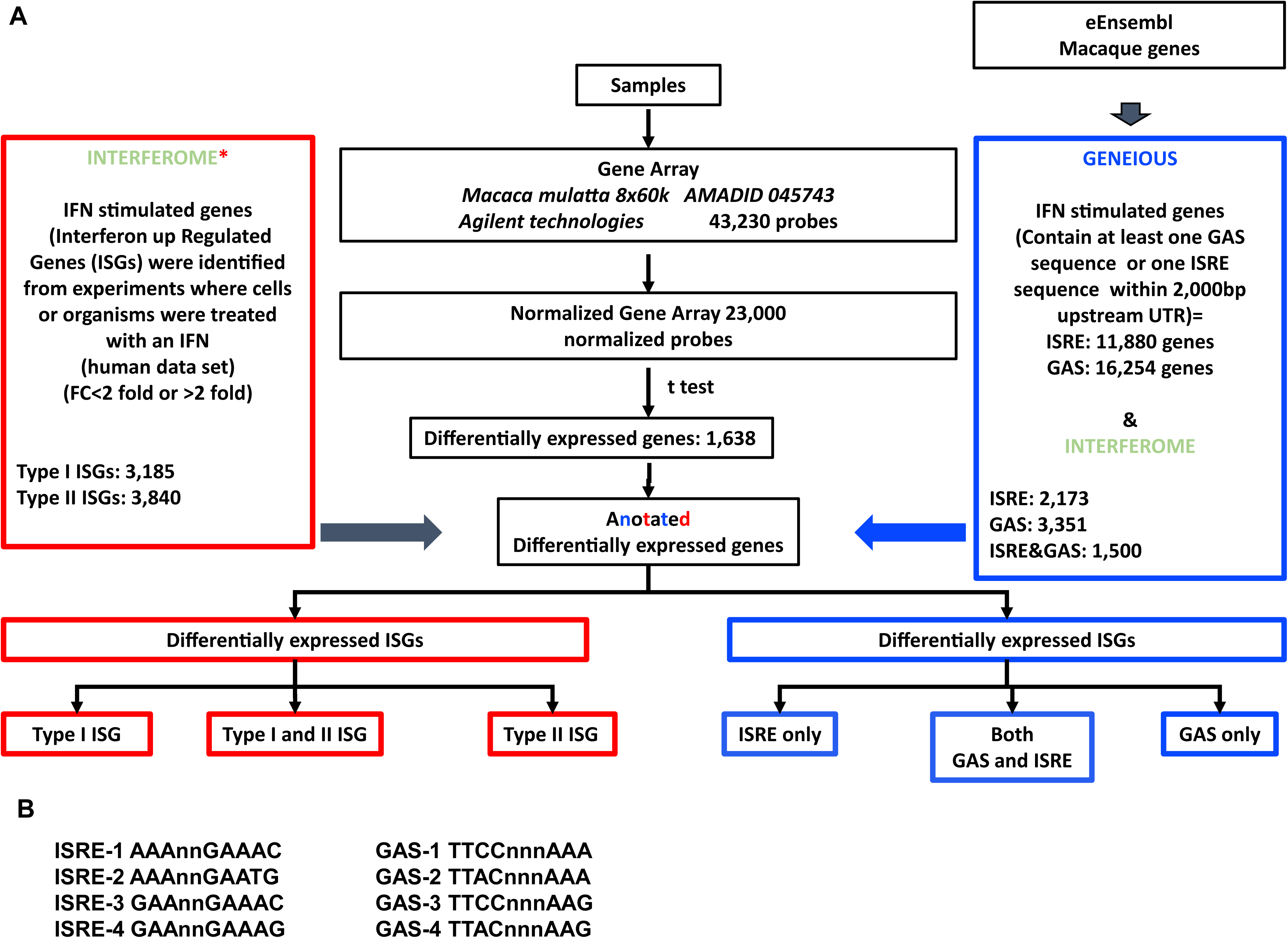
ISG annotations. Differentially expressed interferon-induced genes were annotated using two approaches: the Interferome database helped to identify and classify genes depending on their induction by either type I IFN (α, β), type II IFN (γ),or both IFN types. In addition, Geneious software was used to annotate ISGs depending on the presence of either GAS, ISRE, or both sequence motifs within the 2,000 bp upstream of the 5’UTR of each gene. (**A**) Schematic diagram of annotations. (**B**) ISRE and GAS sequences

**EV Figure 2:**
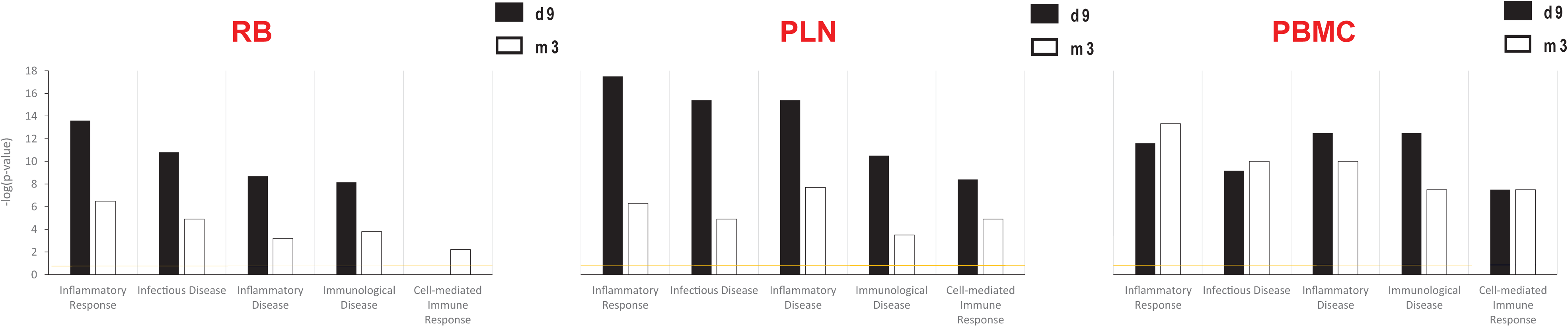
Functional enrichment of differentially expressed genes. IPA enrichment analysis was performed on all differentially expressed genes (non-parametric t-test p < 0.05) independently of their FC. The –log (p-values) of most significant functions are plotted in a histogram format for both D9 p.i. and M3 p.i., in each tissue including **(A)** RBs, **(B)** PLNs, and **(C)** PBMCs.

**EV Figure 3:**
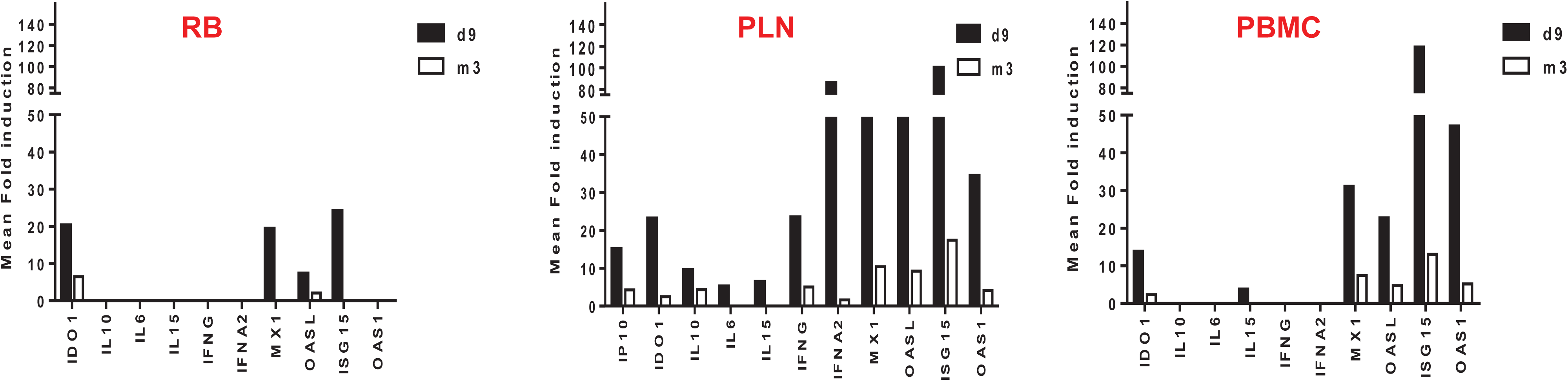
Comparison of the extent of IFNs and selected ISG expression in both phases of infection in RM, PLNs, and PBMCs. The mean FC induction shows that both IFNs and ISGs were more consistently induced in PLNs than in PBMCs or RM.

**EV Figure 4:**
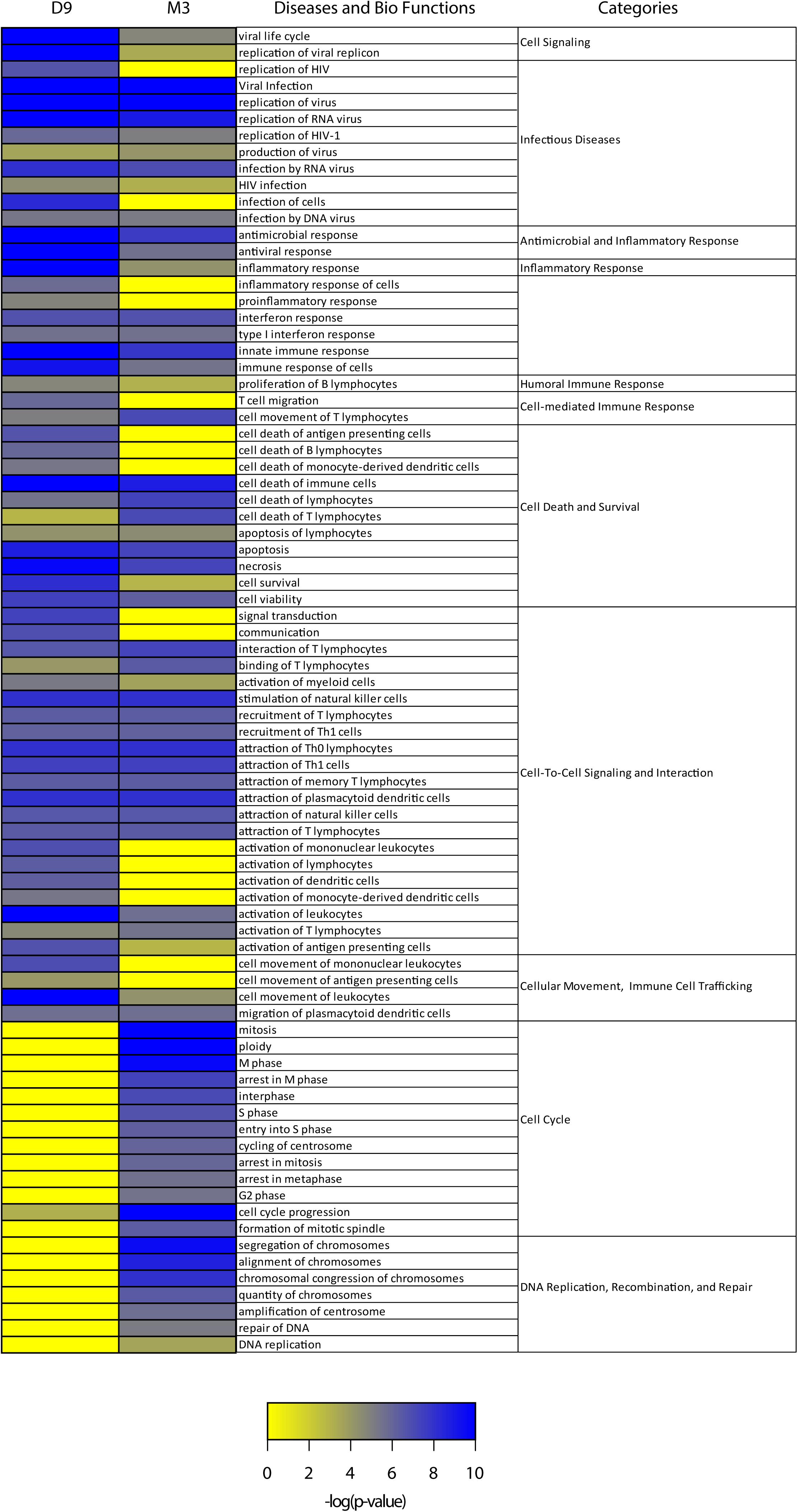
The main IPA functions associated with D9-ISGs and M3-ISGs. IPA enrichment analysis was performed on all ISGs. The –log (p-values) of globally significant functions are plotted in heatmap format for both D9 and M3 p.i.

**EV Figure 5:**
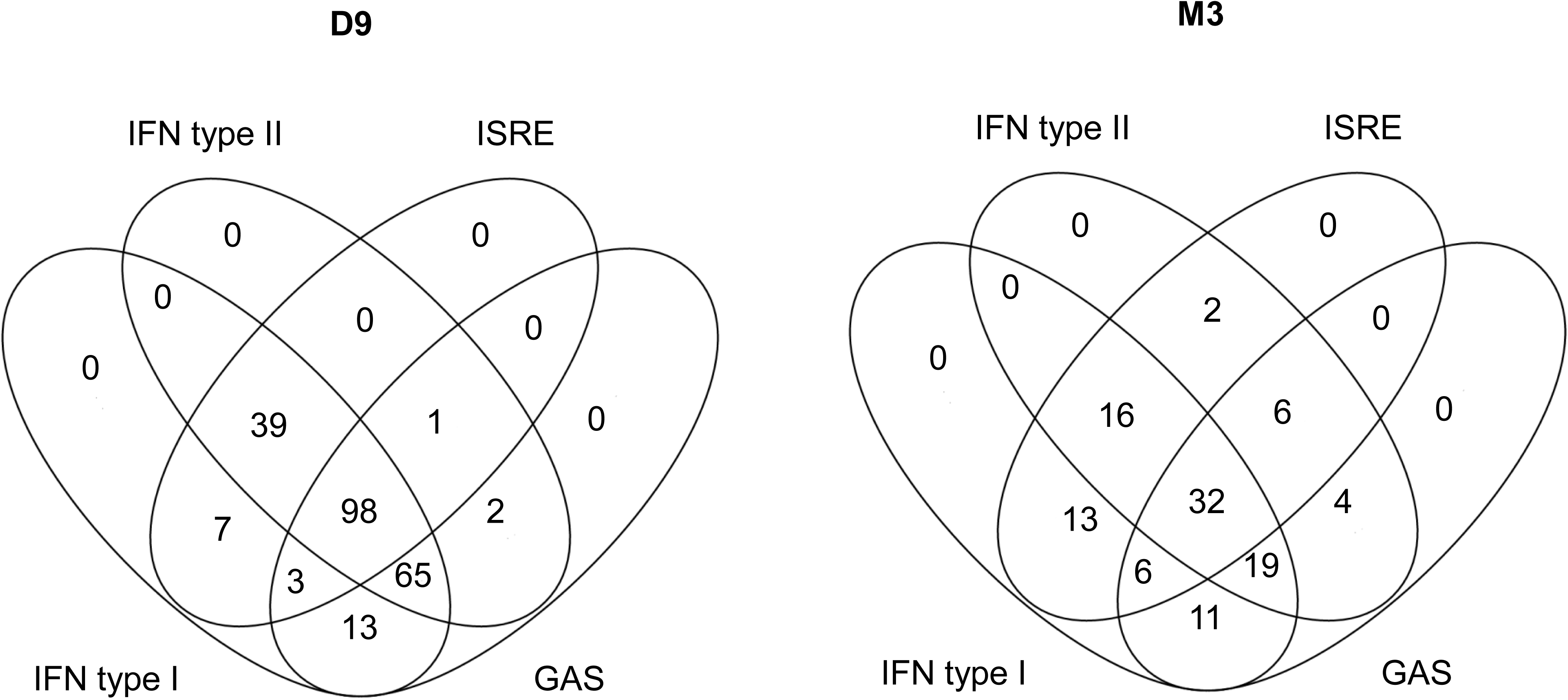
Overlap between Interferome annotation and GAS/ISRE annotation of differentially induced ISGs. Venn diagram of differentially expressed ISGs, showing the number of genes in each annotation subset for D9 or M3 p.i.

